# Natural Variation in Arabidopsi*s* Cvi-0 Accession Uncovers Regulation of Guard Cell CO_2_ Signaling by MPK12

**DOI:** 10.1101/073015

**Authors:** Liina Jakobson, Lauri Vaahtera, Kadri Tõldsepp, Maris Nuhkat, Cun Wang, Yuh-Shuh Wang, Hanna Hõrak, Ervin Valk, Priit Pechter, Yana Sindarovska, Jing Tang, Chuanlei Xiao, Yang Xu, Ulvi Gerst Talas, Maido Remm, Saijaliisa Kangasjärvi, M. Rob G. Roelfsema, Honghong Hu, Jaakko Kangasjärvi, Mart Loog, Julian I. Schroeder, Hannes Kollist, Mikael Brosché

## Abstract

Plant gas exchange is regulated by guard cells that form stomatal pores. Stomatal adjustments are crucial for plant survival; they regulate uptake of CO_2_ for photosynthesis, loss of water and entrance of air pollutants such as ozone. We mapped ozone hypersensitivity, more open stomata and stomatal CO_2_-insensitivity phenotypes of the *Arabidopsis thaliana* accession Cvi-0 to a single amino acid substitution in MAP kinase 12 (MPK12). In parallel we showed that stomatal CO_2_-insensitivity phenotypes of a mutant *cis* (CO_2_-insensitive) were caused by a deletion of *MPK12*. Lack of MPK12 impaired bicarbonate-induced activation of S-type anion channels. We demonstrated that MPK12 interacted with the protein kinase HT1, a central node in guard cell CO_2_ signaling, and that MPK12 can function as an inhibitor of HT1. These data provide a new function for plant MPKs as protein kinase inhibitors and suggest a mechanism through which guard cell CO_2_ signaling controls plant water management.

## Introduction

Human activities have increased the concentrations of CO_2_ and harmful air pollutants such as ozone in the troposphere. During the last 200 years, the CO_2_ concentration has increased from 280 to 400 ppm, and it is predicted to double relative to the preindustrial level by 2050 [1]. Elevated CO_2_ is likely to have complex effects on plant productivity, since CO_2_ is not only a driver of climate change, but also the main substrate for photosynthesis. Altered atmospheric chemistry is not limited to CO_2_, the concentration of tropospheric ozone has more than doubled within the past 100 years [2]. Ozone is a notorious air pollutant causing severe damages to crops; present day global yield reductions caused by ozone range from 8.5-14% for soybean, 3.9-15% for wheat, and 2.2-5.5% for maize [3]. Both of these gases enter the plant through stomata, small pores on the surfaces of plants, which are formed by pairs of guard cells. Guard cells also regulate plant water balance since plants with more open stomata allow faster water evaporation. Water availability is the most limiting factor for agricultural production and can cause vast decreases in crop yields [4]. Thus plants are constantly facing a dilemma; assimilation of CO_2_ requires stomatal opening, but also opens the gates for entrance of harmful air pollutants and leads to excessive water loss. A consequence of increased atmospheric CO_2_ concentration can be higher biomass production [5]; but at the same time plants adjust to elevated CO_2_ by partial closure of stomata [5, 6] and altered developmental program leading to reduced stomatal number [7]. CO_2_-induced stomatal closure reduces water loss, hence it can directly modify plant water use efficiency (WUE) - carbon assimilated through photosynthesis versus water lost through stomata.

Important components of *Arabidopsis thaliana* guard cell CO_2_ signaling are carbonic anhydrases (CA1 and CA4) that convert CO_2_ to bicarbonate and the protein kinase HT1 (HIGH LEAF TEMPERATURE 1) that has been suggested to function as a negative regulator of CO_2_-induced stomatal movements [8, 9]. Ultimately for stomata to close, the signal has to activate protein kinases such as OST1 (OPEN STOMATA1) that in turn activate plasma membrane anion channels, including SLAC1 (SLOW ANION CHANNEL 1) followed by extrusion of ions and water that causes stomatal closure [10-13]. Bicarbonate-induced activation of SLAC1 has been reconstituted in *Xenopus laevis* oocytes [14, 15]. The pathway was shown to consist of RHC1 (RESISTANT TO HIGH CARBON DIOXIDE 1), HT1, OST1 and SLAC1 [14]; while more recently the importance of CA4, aquaporin PIP2;1, OST1 and SLAC1 was demonstrated [15]. Despite that guard cells are perhaps the best characterized single cell signaling system in the plant kingdom, there are still large gaps in our understanding of how CO_2_ signaling in guard cells is regulated, and by which mechanism CO_2_ might regulate plant water management and WUE [5, 16, 17].

Natural variation among *Arabidopsis* accessions provides a rich genetic resource for addressing plant function and adaptation to diverse environmental conditions. The accession Cvi-0 from Cape Verde Islands is extremely sensitive to ozone treatment, has more open stomata than Col-0 and its stomata CO_2_ responses are impaired [18, 19]. A single amino acid change in Cvi-0 MPK12 (MITOGEN-ACTIVATED PROTEIN KINASE 12) was recently shown to affect water use efficiency as well as stomatal size and to impair ABA-induced inhibition of stomatal opening [20]. Together with MPK9, MPK12 regulates stomatal responses to H_2_O_2_, ABA and extracellular Ca^2+^ [21]. MPK12 also regulates auxin signaling in roots [22].

Here we present the results of quantitative trait loci (QTL) mapping and sequencing of near isogenic lines (NILs) of Cvi-0 ozone sensitivity. In a parallel approach we mapped more open stomata and CO_2_-insensitivity phenotypes of a mutant *cis* (CO_2_ *insensitive*). A single amino acid change (G53R) in MPK12 and complete deletion of *MPK12* are the causes of more open stomata and altered CO_2_ responses of Cvi-0 and *cis*, respectively. In kinase activity assays we show that MPK12 can act as an inhibitor of the HT1 kinase, suggesting a mechanism for regulation of stomatal CO_2_ responses.

## Results

### Mapping of Cvi-0 ozone sensitivity phenotypes

Our initial QTL mapping of ozone sensitivity in Cvi-0 placed the two major contributing loci on the lower ends of chromosomes 2 and 3 [18]. To identify the causative loci related to the extreme ozone sensitivity and more open stomata of Cvi-0, we created a near isogenic line (NIL) termed Col-S (for <u>Col</u>-0 ozone <u>s</u>ensitive) through eight generations of backcrossing of Cvi-0 with Col-0 (Fig 1A, S1A Fig and S1 Video). In parallel, ozone tolerance from Col-0 was introgressed to Cvi-0, by six generations of backcrossing, which generated the ozone tolerant Cvi-T (S1 Video). Using these accessions, NILs and recombinant inbred lines (RILs), we mapped the causative ozone QTLs to a region of 90 kb on chromosome 2 and 17.70-18.18 Mbp on chromosome 3 (S1B Fig). We have previously shown that the QTL on chromosome 2 also controls plant water loss and stomatal function [18], we split both QTLs by backcrossing Col-S with Col-0 and obtained the NILs Col-S2 and Col-S3. Both of these were less sensitive to ozone than Col-S (S1A Fig), indicating that these QTLs act additively to regulate ozone sensitivity. Col-S2 (but not Col-S3) showed much higher day-time stomatal conductance than Col-0 (Fig 1B). The mapping resolution on chromosome 3 was not sufficient to identify the causative gene. Hence, we focused on Col-S2 and its role in stomatal function. Within the 90 kb mapping region on chromosome 2 one gene, At2g46070 encoding a MAP kinase MPK12, shows strong preferential guard cell expression [21]. A single point mutation was found in Cvi-0 *MPK12* leading to a glycine to arginine substitution at position 53 of the protein.

**Figure 1.**
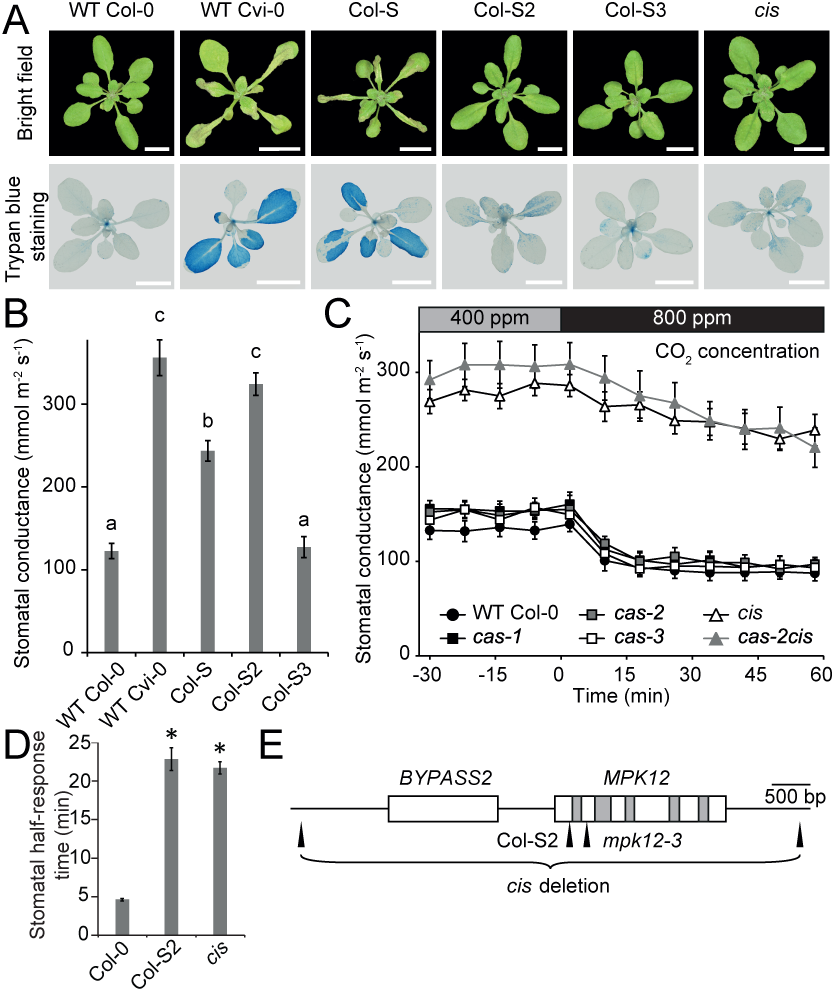
Mapping of Cvi-0 ozone sensitivity and Cvi-0 and *cis* stomatal responses. (**A**) Tissue damage after 6 h of O_3_ exposure (350 ppb). Visual damage of plant rosettes (upper images) and cell death visualized with trypan blue staining (lower images). Scale bar 1 cm. (**B**) Stomatal conductance of Col-0, Cvi-0 and NILs (mean ± SEM, n = 7-12. (**C**) Elevated CO_2_ (800 ppm) induced stomatal closure in intact whole plants (n = 9-10, except *cas-2* n = 3. Experiment was repeated at least three times with similar results. (**D**) Stomatal half-response times to elevated CO_2_ (800 ppm). Error bars indicate ± SEM (n = 13). Pooled data from two experimental series are shown. **(E)** Gene model of *MPK12*(At2g46070) and *BYPASS2* (AT2g46080). The deletion mutant *cis* (renamed as *mpk12-4*) has a 4772 bp deletion (end and start indicated). Col-S2 has a G to C missense mutation at position 157 of *MPK12*, which leads to G53R substitution in MPK12. The *mpk12-3* mutant has a SAIL T-DNA insertion site in the second exon of *MPK12*. White boxes refer to exons, grey boxes to introns and black line to intergenic regions. Small letters (B) and asterisks (D) denote statistically significant differences according to 1-way ANOVA with Tukey HSD *post hoc* test.

### Stomata-related phenotypes of Cvi-0 and *cis* are caused by mutations in *MPK12*

In a parallel project we observed phenotypic discrepancy between different alleles of *cas* (Calcium-sensing receptor [23]). The *cas-2* (GABI-665G12) line had more open stomata and impaired CO_2_ responses, whereas this was neither observed in *cas-1* nor in *cas-3* (Fig 1C and S1C Fig). In a backcross with Col-0 the T-DNA insert in *cas-2* was removed, thereby generating the mutant *cis* (CO_2_ *insensitive*). Both *cis* and Col-S2 had impaired responses to high CO_2_ (800 ppm) leading to longer half-response times, but a residual CO_2_ response could still be observed (Fig 1D and S1E Fig).

Mapping and whole genome sequencing of *cis* × C24 populations revealed a complete deletion of the *MPK12* gene and its neighbor *BYPASS2* in *cis* (Fig 1E and S1D Fig). Thus, *cis* was renamed to *mpk12-4.* A second mutant (*gdsl3-1*) from the GABI collection (GABI-492D11) contained an identical deletion of *BYPASS2* and *MPK12* (S2 Fig). We also identified a line with a T-DNA insert in exon 2 of *MPK12* from the SAIL collection (Fig 1E), which was recently named *mpk12-3* [24]. No full-length transcript was found in *mpk12-3* (S3 Fig). SALK T-DNA insertion lines of *MPK12* were previously described as lethal [21, 22]; similarly we were unable to retrieve homozygous plants of the same alleles possibly indicating the presence of an additional T-DNA in an essential gene. The new *mpk12* deletion, SAIL insertion and Col-S2 point mutation alleles allowed a detailed characterization of the role of MPK12 in stomatal regulation.

Stomatal conductance was higher throughout the day in all three lines: Col-S2, *mpk12-3* and *mpk12-4* (Fig 2A), suggesting that the amino acid substitution in Cvi-0 MPK12 leads to loss of function. Furthermore, Col-0 transformed with *MPK12* from Cvi-0 and F1 plants from a cross of Col-S2 and Col-*gl1* showed stomatal conductance similar to Col-0, excluding the option that the G53R substitution in MPK12 would lead to gain of function (S1F-S1G Fig). Increased stomatal conductance may result from an increased number of stomata, larger stomata or more open stomata. However, the stomatal index, length and density did not differ between the lines, indicating that MPK12 regulates a function related to the stomatal aperture (S4 Fig). Because of the higher degree of stomatal opening, the instantaneous WUE was decreased for *mpk12-3*, *mpk12-4* and Col-S2 (Fig 2B). Altered WUE was previously also seen in *mpk12-1* and a NIL with Cvi-0 MPK12 in L*er* [20]. Cvi-0 and Col-S2 were complemented by expression of MPK12 from Col-0 (Fig 2C and 2D). Similarly, *mpk12-4* was complemented by expression of Col-0 MPK12 but not by Cvi-0 MPK12 (Fig 2E). We conclude that MPK12 is a crucial regulator of stomatal conductance and a single amino acid substitution (G53R) in Cvi-0 leads to a loss of function in MPK12.

**Figure 2.**
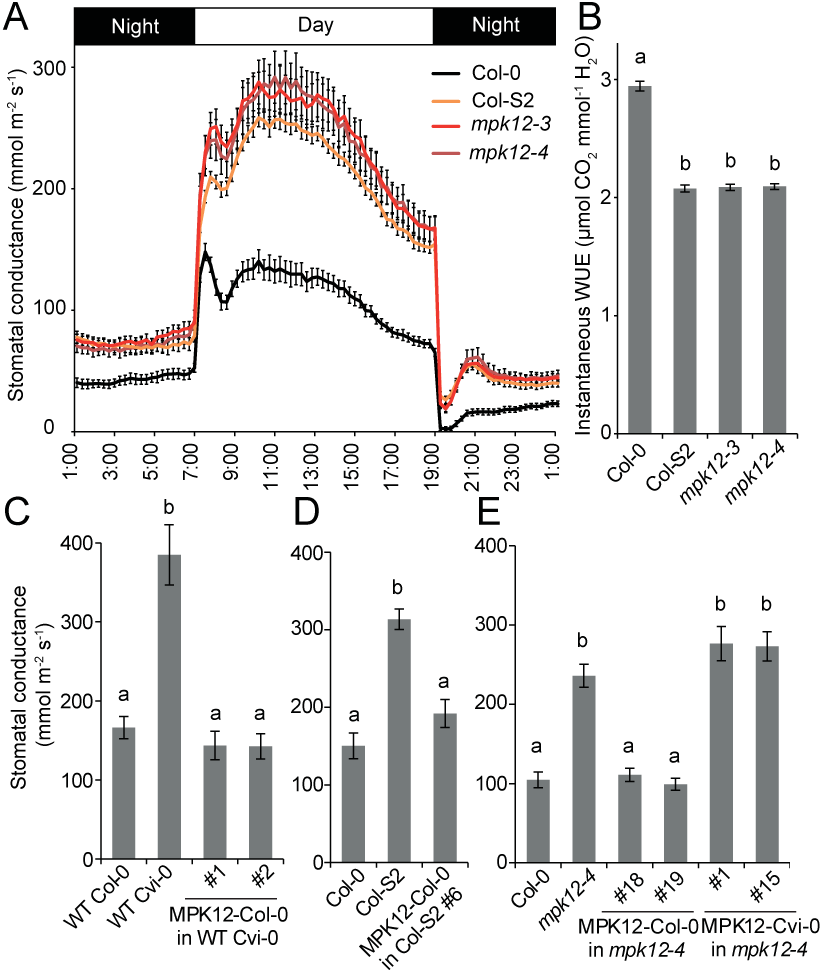
Stomatal conductance of the NIL Col-S2, *mpk12* mutants and complementation lines. (**A**) Diurnal pattern of stomatal conductance with 12h/12h light-dark periods (n = 13-16). (**B**) Instantaneous water use efficiency (WUE) measured as an average of day-time light period from 09:00 to 17:00 (n = 3-16). (**C**) Stomatal conductance of Cvi-0 transformed with Col-0 *MPK12* and its promoter in T2 generation (n = 9). (**D**) Stomatal conductance of Col-S2 complementation line in T2 generation transformed with Col-0 *MPK12* and its promoter (n = 5-8). (**E**) Stomatal conductance of T3 transformants in the *mpk12-4* background transformed with either the Col-0 or Cvi-0 version of MPK12 and its respective promoter (n = 5-6). All graphs present mean ± SEM. Small letters denote statistically significant differences according to 1-way ANOVA with Tukey HSD *post hoc* test for either unequal (B, D, E) or equal sample size (C).

### MPK12 functions in guard cell CO_2_ signaling

Reduction of CO_2_ levels inside the leaf [25] is a signal that indicates shortage of substrate for photosynthesis and triggers stomatal opening. The rate of stomatal opening in response to low CO_2_ was severely impaired in *mpk12* and Col-S2 (Fig 3A and S5A Fig). Another signal for stomatal opening is light; this response was intact in *mpk12* (S5B-C Fig). The hormone abscisic acid (ABA) has dual roles in stomatal regulation; it induces stomatal closure, but also inhibits light-induced stomatal opening. The latter response was impaired in *mpk12* and Col-S2 (Fig 3B and S5C Fig). Stomata close in response to several signals including darkness, reduced air humidity, ozone pulse, elevated CO_2_ and ABA. Of these, only the response to elevated CO_2_ was impaired in *mpk12* and Col-S2 (Fig 3C, 3D, S5D-H and S6 Fig). CO_2_ signaling is impaired in the carbonic anhydrase double mutant *ca1 ca4* [9] and the product of carbonic anhydrase, bicarbonate, activates S-type anion currents [11]. In Col-S2 and *mpk12-4,* bicarbonate-induced S-type anion currents were strongly impaired (Fig 3E). Collectively, these data indicated that MPK12 has an important role in the regulation of CO_2_-induced stomatal movements in *Arabidopsis*.

**Figure 3.**
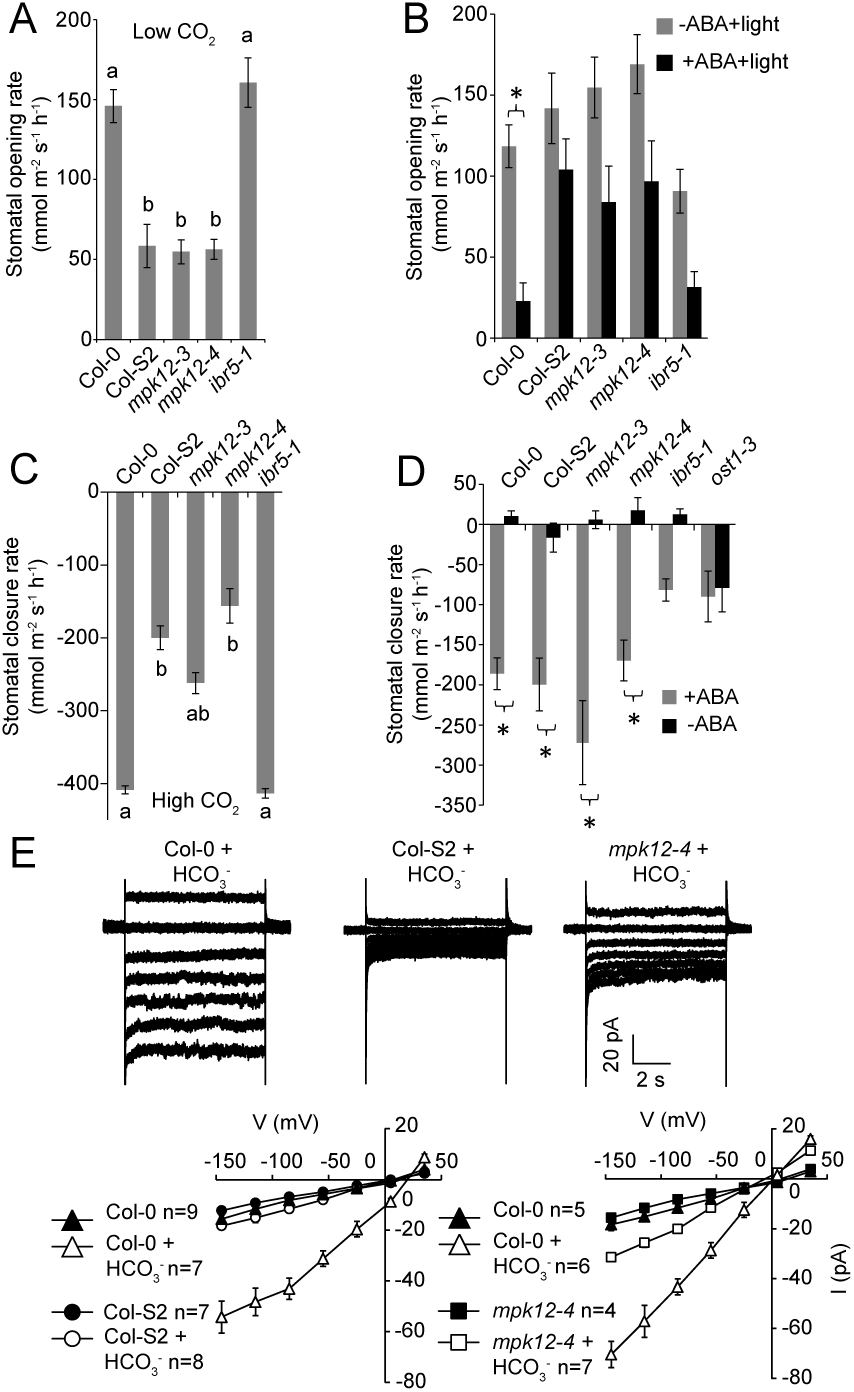
Stomatal responsiveness of the NIL Col-S2 and *mpk12* mutants to opening and closing stimuli. (**A**) Stomatal opening rate induced by 100 ppm CO_2_ in whole plants (58 min after induction; n = 12-13). (**B** Light-induced stomatal inhibited by 2.5 μM ABA in whole plants (24 min after induction; n = 16-18). (**C**) Stomatal closure rate induced by 800 ppm CO_2_ in whole plants (10 min after induction; n = 12-13). (**D**) Stomatal closure rate induced by spraying whole plants with 5 μM ABA solution (24 min after induction n = 12-14). (**E**) Slow type anion channel activity in guard cell protoplasts treated with HCO_3_^-^ or mock treatment. Small letters (A, C) and asterisks (B, D) indicate statistically significant differences according to 1-way ANOVA and 2-way ANOVA with Tukey HSD unequal N *post hoc* tests (p < 0.05), respectively. Error bars mark ± SEM.

### MPK12 interacts with the protein kinase HT1

Only a few regulators of stomatal CO _2_signaling in *Arabidopsis* have been identified. These include the protein kinases HT1 and OST1 [8, 11, 12]. To find the interaction partners of MPK12 we conducted pair-wise split-ubiquitin yeast two-hybrid (Y2H) assays against several kinases and phosphatases involved in stomatal signaling, and against two MPK phosphatases that regulate MPK activity, IBR5 (INDOLE-3-BUTYRIC ACID RESPONSE 5) and MKP2 (MAPK PHOSPHATASE 2) (Fig 4A, 4B and S7A-B Fig). A strong interaction was observed between MPK12 and HT1 in yeast, but not for any of the other kinases or phosphatases except for MPK12 and IBR5. The MPK12-HT1 interaction was also confirmed in *Nicotiana benthamiana* with bimolecular fluorescence complementation (BiFC) (Fig 4C) and split luciferase complementation assay (S7C Fig). Strong interaction between MPK12 and HT1 was observed in the cell periphery (Fig 4C). Recently HT1 was shown to be a plasma membrane associated protein [26]. In contrast, Col-0 and Cvi-0 MPK12-YFP were located inside the cell (S8A-D Fig), hence it is likely that the interaction with HT1 brings MPK12 to the plasma membrane. HT1 interacted with both Col-0 and Cvi-0 version of MPK12, however, HT1 showed weaker interaction with Cvi-0 MPK12 (G53R) both in BiFC and quantitative Y2H assays (Fig 4B and 4C). MPK11, an MPK from the same group as MPK12 [27], did not interact with HT1 (Fig 4C).

**Figure 4.**
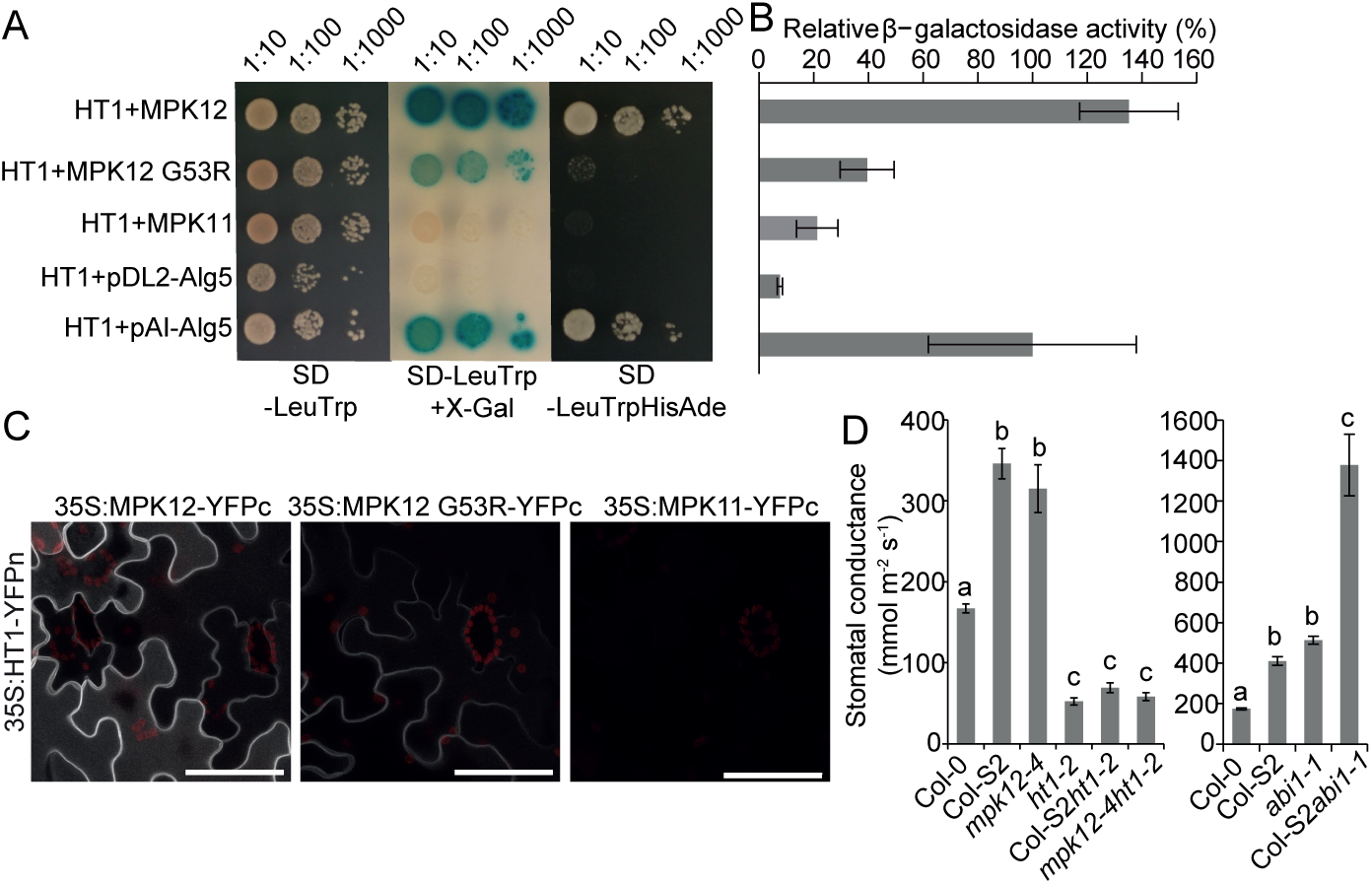
MPK12 interacts with HT1. (**A**) Split-ubiquitin yeast two-hybrid assay on the SD-LeuTrp plate (left and middle panels) for the presence of both bait and prey plasmids; X-gal overlay assay (middle) and growth assay on the SD-LeuTrpHisAde plate (right) show HT1 interaction with MPK12, similar to the positive control (pAI-Alg5). Only weak or no interaction was detected with MPK12 G53R or MPK11, similar to the negative control (pDL2-Alg5). (**B** Quantitative β-galactosidase assay from pools of 10 colonies each. Activities are shown as the percentage of the positive control (± SEM, n = 3). (**C**) BiFC images from the same infiltrated tobacco (*N. benthamiana*) leaf with identical confocal microscopy acquisition settings. Scale bar = 50 μm. **D**) Steady-state stomatal conductance of Col-S2 *ht1-2*, *mpk12-4 ht1-2* and Col-S2 *abi1-1* double mutants (mean ± SEM, n = 11-13; 1-way ANOVA, Tukey HSD *post hoc* test for unequal sample size). Experiments were repeated at least three times.

IBR5 interacts with MPK12 and regulates auxin signaling in roots [22]. If IBR5 would regulate the activity of MPK12 also in stomatal CO_2_ signaling, the *ibr5* mutant would have been expected to display altered CO_2-_ related stomatal phenotypes. However, *ibr5-1* exhibited wild type stomatal phenotypes in response to CO_2_ changes (Fig 3A, 3C, S5A and S5E Fig), indicating that IBR5 does not play a role in the regulation of MPK12 activity in the guard cells.

### MPK12 inhibits HT1 activity

The position of MPK12 in ABA and CO_2_ signaling was further explored through genetic analysis. The Col-S2 *ht1-2* and *mpk12-4 ht1-2* double mutants had closed stomata similar to *ht1-2* (Fig 4D), suggesting that *HT1* is epistatic to *MPK12*. The strong impairment of stomatal function in *abi1*-*1* (*ABA insensitive1-1*) was additive to Col-S2 in the double mutant Col-S2 *abi1-1* (Fig 4D). Hence, signaling through MPK12 seems to act, at least to some extent, independently of the core ABA signaling pathway. Taken together, the MPK12-HT1 interaction (Fig 4A-C) and the epistasis between *ht1-2* and *mpk12-4* (Fig 4D) suggest that MPK12 regulates the activity of HT1. To test this directly, we performed *in vitro* kinase assays with casein as the substrate for HT1 (Fig 5A). HT1 displayed strong autophosphorylation and phosphorylated casein efficiently. Addition of the Col-0 version of MPK12 and its hyperactive version (Y122C) effectively inhibited HT1 activity (Fig 5A and quantified in Fig 5B). A point mutated version (K70R) designed to remove the kinase activity of MPK12 also inhibited autophosphorylation activity of HT1 and phosphorylation of casein by HT1, although to a lesser extent (Fig 5B). Importantly, the Cvi-0 version of MPK12 (G53R) displayed strongly suppressed inhibition of HT1 activity (Fig 5A, and 5B). MPK12 did not phosphorylate the kinase dead version of HT1 (K113M) (Fig 5C). Kinase dead HT1 (K113M) was used as substrate, since the strong autophosphorylation activity of HT1 would otherwise have obscured the result. Wild type MPK12 and hyperactive Y122C displayed autophosphorylation, whereas MPK12 (G53R) as well as MPK12 (K70R) had lost the autophosphorylation activity indicating that the G53R substitution in Cvi-0 MPK12 disrupts the kinase activity of the protein (Fig 5C). Modeling of the MPK12 structure supported a major role for G53, since it was located in the glycine rich loop important for binding of ATP (S9 Fig). MPK11 that belongs to the same group as MPK12, was not able to affect HT1 kinase activity indicating specificity for MPK12 in the inhibition of HT1 (S10 Fig).

**Figure 5.**
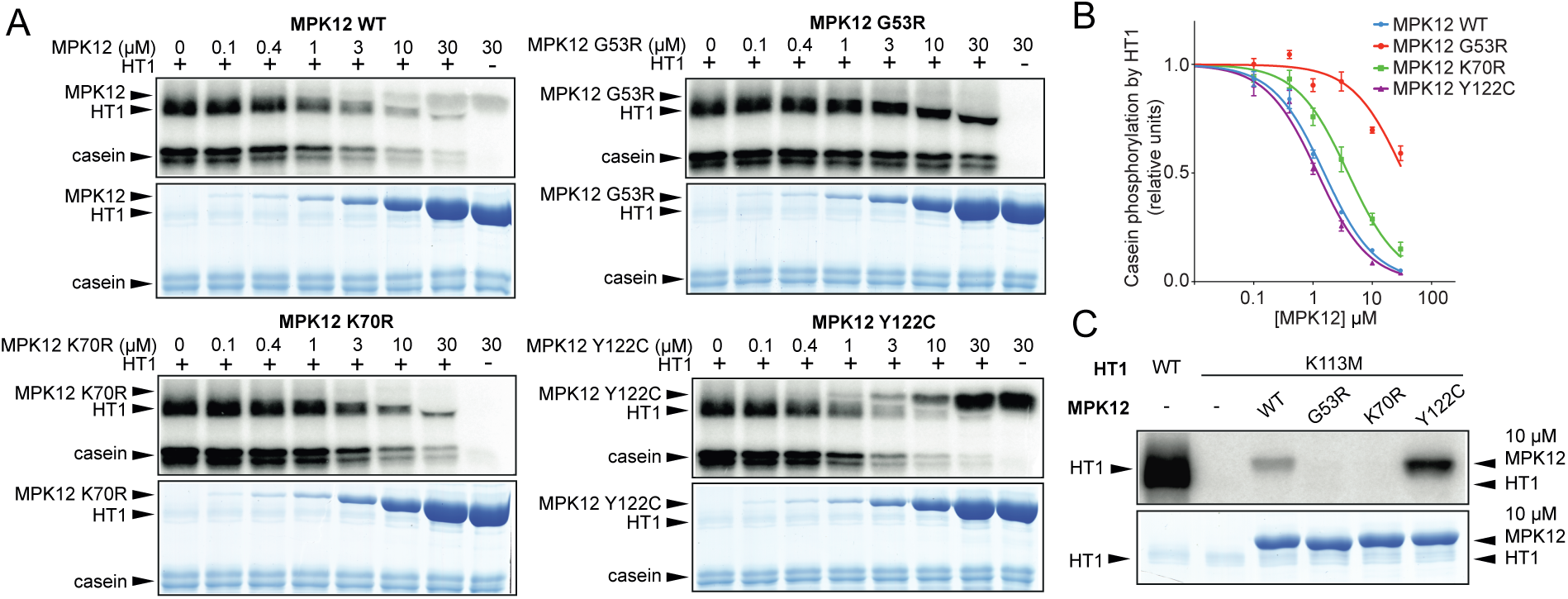
Regulation of HT1 by MPK12. (**A**) *In vitro* HT1 kinase activity inhibited by different versions of MPK12 (upper panel: autoradiography of the SDS PAGE gel; lower panel: Coomassie stained SDS PAGE). Reaction mixture was incubated 30 min. (**B**) Casein phosphorylation by HT1 with different MPK12 concentrations (mean ± SEM, n = 3) (**C**) Kinase dead HT1 K113M was not *in vitro* phosphorylated by different versions of MPK12.

We conclude that the stomatal phenotypes of *mpk12* mutants and Cvi-0 can be explained by a lack of inhibition of HT1 activity by MPK12, which leads to more open stomata and impaired CO_2_ responses (Fig 2A, 3, 5A, 5C and 6).

**Figure 6.**
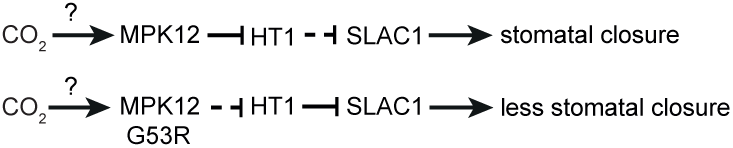
Schematic representation of guard cell CO_2_ signaling pathway via MPK12 and HT1. The CO_2_ signal to stomatal closure is transmitted through MPK12, which inhibits the negative regulator HT1. Cvi-0 MPK12 G53R cannot inhibit HT1.

### Evolutionary considerations of MPKs in the regulation of guard cell CO_2_ signaling

MPK12 belongs to the same group of MPKs as MPK4, a crucial regulator of pathogen and stress responses [27]. In tobacco, the silencing of MPK4 impaired CO_2_-induced stomatal closure [28]. This raises the question whether: (I) MPK4 and MPK12 have a shared function in the regulation of stomatal responses in *Arabidopsis*; or (II) they have specialized function, for example through gene duplication, leading to MPK4 that regulates pathogen responses and MPK12 that regulates stomatal responses to CO_2_. If Arabidopsis MPK12 is derived from MPK4 then this should be visible in a phylogenetic analysis. We identified all MPK sequences from representative angiosperm species and performed a phylogenetic analysis (S11 Fig). The gene duplication event giving rise to MPK12 from ancestral MPK4 appeared to be specific to *Brassicaceae*. The Arabidopsis *mpk4* mutant is severely dwarfed preventing accurate stomatal measurements. However, guard cell specific silencing of *MPK4* in *mpk12-4* led to stronger stomatal CO_2_ insensitivity (Hõrak et al. submitted), hence in guard cells MPK4 is acting redundantly with MPK12 in stomatal CO_2_ signaling. The phylogeny analysis also indicated that *MPK4* was independently duplicated in both grasses and *Solanaceae* (S11 Fig), thus it is likely that in these species several MPKs related to MPK4 may regulate CO_2_–induced stomatal responses. However, since MPK11 did not inhibit HT1 activity (S10 Fig), the function of MPKs as kinase inhibitors may be restricted to MPK12 and its closest relatives including MPK4.

## Discussion

Natural variation within a species holds great potential to identify regulatory mechanisms that are not easily uncovered through mutant screens. The Cvi-0 accession originates from the Southern border of the *Arabidopsis* distribution area, the Cape Verde Islands. The Ler × Cvi RIL population was one of the first RILs produced and it has been phenotyped for multiple traits [29]. Despite this, only a few QTLs from Cvi-0 have been identified at the molecular level. Our earlier research identified a locus related to ozone sensitivity and more open stomata phenotype of Cvi-0 in chromosome 2 [18]. Recently the G53R substitution in MPK12 that affects plant water use efficiency was identified by using the Ler × Cvi populations, but the biochemical function of MPK12 in stomatal regulation was not further investigated [20]. Here, we generated NILs by backcrossing Cvi-0 eight times to Col-0 and show that the same natural mutation in Cvi-0 and lack of MPK12 in *cis* are the causes of ozone sensitivity, more open stomata and altered CO_2_ responses of *Arabidopsis* plants. Furthermore, we identify a mechanism by which MPK12 regulates the activity of the protein kinase HT1, a negative regulator of CO_2_-induced stomatal movements. The regulators of HT1 have remained largely unknown, despite the exceptionally strong CO_2_-insensitivity phenotype of plants with impaired HT1 function [8, 11]. Our findings provide the first evidence for the role of MPK12 in guard cell CO_2_ signaling and how this relates to plant water management.

The role of MPKs in *Arabidopsis* guard cell signaling has concentrated on MPK9 and MPK12, which are preferentially expressed in guard cells. Plants with point mutations in *MPK9* (*mpk9-1,* L295F) and *MPK12* (*mpk12-1,* T220I) had wild type ABA responses, but *mpk12-1* has decreased WUE [20]. The *mpk9-1, mpk12-1* and *mpk12-2* alleles are TILLING (Targeting Induced Local Lesions in Genomes) lines in Col-*erecta* background and the previously characterized MPK12-Cvi NIL is in L*er* background [20, 21]. Mutations in *ERECTA* modifies transpiration efficiency and stomatal density, which may have influenced some of the previously described *mpk12-1* phenotypes [20, 30]. In contrast, the full knockout alleles described here, *mpk12-3* and *mpk12-4*, are in Col-0 and imply a major function for MPK12 in CO_2_ signaling. Additional roles for MPK12 in stomatal responses have been inferred through the use of the double mutant *mpk9-1 mpk12-1* that has impaired stomatal closure responses to ABA and H_2_O_2_ treatment, and impaired S-type anion channel activation in response to ABA and Ca^2+^ [21]. It is also highly susceptible to *Pseudomonas syringae* infection and impaired in yeast elicitor, chitosan and methyl jasmonate induced stomatal closure [31]. Since the *mpk9 mpk12* double mutant appears to be more severely impaired in abiotic and biotic stomatal responses and S-type anion channel activation than the loss of function MPK12 alleles (Fig 3), it is possible that MPK12 together with MPK9 regulates stomatal aperture in response to various signals. MPK12 also regulates auxin responses in the root [22, 24]. However, beyond the observation that plants with impaired MPK12 are hypersensitive to auxin inhibition of root growth, no details about the targets of MPK12 in roots are known.

HT1 is a negative regulator of CO_2_ signaling and the *ht1-2* mutant has more closed stomata displaying constitutive high CO_2_ response at ambient CO_2_ levels (Fig 4D, [8]). The opposite phenotypes of *mpk12* and *ht1-2* allowed us to use genetic analysis to position MPK12 in the guard cell signaling network. The stomata of *mpk12 ht1-2* were closed, thus positioning MPK12 up-stream of HT1 and possibly as a direct regulator of HT1 (Fig 4D). CO_2_ signaling in guard cells is initiated through the production of bicarbonate by carbonic anhydrases and bicarbonate initiates signaling leading to activation of S-type anion channels [9, 11]. In *mpk12*, the bicarbonate-dependent activation of S-type channels was impaired as previously found for the plants with impaired OST1 and SLAC1 (Fig 3E; [11]). The combined evidence from *mpk12* phenotypes, genetic analysis, and measurements of S-type anion currents all pointed towards MPK12 as a crucial regulator of CO_2_ signaling acting through HT1. Indeed, HT1 kinase activity was inhibited in the presence of Col-0 MPK12 but not by the Cvi-0 version of MPK12 (Fig 5A). Thus, the inhibitory function of MPK12 was impaired by the G53R amino acid substitution, probably by its weaker interaction with HT1 (Fig 4 and 5A). This explains the similar phenotypes of the NIL Col-S2, *mpk12-3* and *mpk12-4*; they all display lack of inhibition of the negative regulator HT1 leading to higher conductance at ambient CO_2_ levels. Further support for the regulatory interplay between HT1 and MPK12 is provided by the isolation of a dominant mutation in *HT1*, *ht1*-*8D*, which opposite to *ht1-2,* has constitutively more open stomata and is biochemically resistant to inhibition by MPK12 (Hõrak et al. submitted).

Recently two independent studies used *Xenopus laevis* oocytes as a heterologous expression system to reconstitute bicarbonate-induced activation of the SLAC1 anion channel [14, 15]. Tian et al., [14] reported that a MATE-type transporter, RHC1 functions as bicarbonate-sensing component that inactivates HT1 and promotes SLAC1 activation by OST1. More recently it was demonstrated that expression of RHC1 alone was sufficient to activate ion currents in oocytes; these currents were independent of bicarbonate, calling into question the role of RHC1 as a bicarbonate sensor [15]. Furthermore, it was shown that SLAC1 activation can be reconstituted by extracellular bicarbonate in the presence of aquaporin PIP2.1, CA4 and the protein kinases OST1, CPK6 and CPK23 [15]. However, in the guard cell, any proposed CO_2_ signaling pathway should include HT1, since plants with mutations in HT1 completely lack CO_2_–induced stomatal responses [8]. We showed that bicarbonate-induced S-type anion currents were strongly impaired in guard cells protoplasts, which lacked functional MPK12 (Fig 3E). Thus, MPK12 and possibly other MPKs that are expressed in guard cells play a role in controlling the activity of HT1 and future research should identify the signaling pathway upstream of MPK12. Dissection of different domains in SLAC1 revealed that the CO_2_ signal may involve the transmembrane region of SLAC1, whereas ABA activation of SLAC1 requires intact N- and C-terminus [32]. Hence, ABA and CO_2_ regulation of SLAC1 could use different signaling pathways, and this may explain the lack of strong ABA phenotypes in plants carrying mutations in *MPK12*.

Stomata also regulate other aspects of leaf function, for example pathogen entry into leaves. Cvi-0 has altered phenotypes in many traits including drought and pathogen resistance [29, 33, 34]. All of these traits are at least partly regulated through proper stomatal function, thus the MPK12-HT1 regulatory module identified here may influence many of the previously observed phenotypes of Cvi-0. G53 is conserved in all *Arabidopsis* MPKs [20], and is located in the glycine-rich loop that coordinates the gamma-phosphate of ATP. Since the G53 of MPK12 is replaced with the bulky and charged arginine in Cvi-0, it would be expected to impair kinase activity, which was indeed observed (Fig 5C). R53 of Cvi-0 MPK12 is predicted to be positioned at the protein surface, which could explain the weaker interaction with HT1 (Fig 4 and S9 Fig). We propose that negative regulation of stomatal CO_2_ signaling by HT1 is alleviated through interaction with MPK12 and MPK4 (Fig 6; Hõrak et al. submitted). The similar impairment of CO_2_ signaling and activation of S-type anion channels when MPK4 function is silenced in tobacco [28] and when MPK12 is absent in *Arabidopsis* (Fig 3E), combined with the phylogenetic analysis showing that MPK4 has duplicated also in grasses (S11 Fig) indicate that the MPK12-HT1 CO_2_ regulatory module identified here may be broadly conserved throughout angiosperms. Further studies into the mechanisms controlling activation of MPKs in guard cells will help to identify the early events in the perception of altered CO_2_ concentrations.

## Materials and methods

### Plant Material and Growth Conditions

Col-0, Col-*gl*, Cvi-0, *gdsl3-1* (GABI-492D11; CS447183), *cas-1* (SALK_070416), *cas-2* (GABI-665G12) and *cas-3* (SAIL_1157_C10) were from the European Arabidopsis Stock Centre (www.arabidopsis.info). Seeds of *ht1-2* were a gift from Dr. Koh Iba. Col-0 × Cvi-0 RILs were obtained from INRA Versailles. The *abi1-1* allele used was in Col-0 accession. Double mutants and other crosses were made through standard techniques and genotyped with PCR based markers (S1 Table).

For ozone screening, seeds were sown at high density on a 1:1 v/v mixture of vermiculite and peat (type B2, Kekkilä, Finland), and kept for 2 d at 4 °C for stratification. The plants were grown in controlled growth chambers (Bio 1300,Weiss Umwelttechnik, Germany) under a 12 h photoperiod, 23/19 °C day/night temperature and 70/90% relative humidity or growth rooms with equivalent growth conditions. The average photosynthetic photon flux density (PPFD) during the light period was 200 mmol m^−2^ s^−1^. When seedlings were 1 week old, they were transplanted into 8 × 8 cm pots at a density of five plants per pot.

Three-week-old plants were exposed to ozone in growth chambers under the same conditions as they were grown until the experiments. Ozone exposure was acute (300– 350 ppb for 6 h) and started 2 h after light was switched on. Ozone damage was visualized with trypan blue stain or quantified as electrolyte leakage.

### Mapping of Cvi-0 ozone sensitivity QTLs

Near isogenic lines (NILs) were created by crossing Col-0 with Cvi-0 and selecting the most ozone sensitive plant in F2 and backcrossing to Col-0 for eight generations (generating Col-S) or selecting the most tolerant plant and backcrossing to Cvi-0 for six generations (generating Cvi-T). The genomes of Cvi-0 and Col-S were sequenced at BGI Tech Solutions (Hongkong) with Illumina technology and the genomes of Col-S and Cvi-T were sequenced at the DNA Sequencing and Genomics lab, University of Helsinki, with SOLiD technology. The 90 bp long Illumina paired end sequencing library reads were mapped onto the Col-0 reference genome (TAIR10) with using Bowtie2 aligner (version 2.0.0-beta7;[35]) in “end-to-end” alignment mode, yielding an average genomic sequence coverage of 45 fold. Variation calling and haplotype phasing was performed with the help of samtools (Tools for alignments in the SAM format, Version: 0.1.18; [36]). Based on the aligned sequences various PCR-based markers (S1 Table) were designed to genotype Cvi-0 versus Col-0 in the NILs and informative RILs from the INRA Versailles Col-0 × Cvi-0 RIL population. The markers were also used to genotype ozone sensitive individuals from segregating F2 populations.

### Mapping of *cis* mutation

Mapping population was created by crossing *cis* (Col-0) and C24 as an *Arabidopsis* genotype with low stomatal conductance. High water loss from excised leaves and decreased responses to high CO_2_ were used as a selective trait. Rough mapping with 22 markers using 59 F2 samples showed linkage to the bottom of chromosome 2, at the marker UPSC_2-18415 at 18,4 Mbp. Pooled genomic DNA from 66 selected F3 lines was used for sequencing. Whole genome sequencing was conducted with Illumina HiSeq 2000 and the reads were mapped against Col-0 genome (release TAIR10) by BGI Tech Solutions (Hongkong). For mapping the genomic area of the mutation the Next Generation Mapping (NGM) tool was used [37], which positioned the mutation on chromosome 2 between 18 703 644-19 136 098 bp. The deletion mutation in *cis* was verified by PCR to be 4 770 bp (at the position 18 945 427-18 950 196 bp).

### Complementation lines

*MPK12* and its promoter were amplified from Col-0 or Cvi-0 genomic DNA using Phusion (Thermo Scientific) and Gateway (Invitrogen) cloned into entry vector pDONR-Zeo. Subsequently, the genes were cloned into pGWB13 and pMCD100. Plants were transformed with floral dipping [38].

### Southern blotting analyses

Total DNAs from different genotyping plants were extracted by CTAB method, and 12 micrograms of total DNA was digested by Hind III or EcoRI. The DNAs were running on the gel and transformed onto Nylon membrane. Hybridization was performed with digoxigenin-labeled specific genomic DNA amplified by primers F3 and R4 for 12 h. The membrane was washed several times by washing buffer and Maleic acid buffer. The membrane was blocked by blocking solution for 1 h at room temperature and washed and incubated with anti-DIG-AP for 30 min. Detection was performed using substrate DIG CSPD.

### Plant growth and experimental settings for gas-exchange measurements

Seeds were planted on soil mixture consisting of 2:1 (v:v) peat:vermiculite and grown through a hole in a glass plate covering the pot as described previously (34). Plants were grown in growth chambers (MCA1600, Snijders Scientific, Drogenbos, Belgium) at 12/12 h day/night cycle, 23°/20° C temperature, 100 µmol m^−2^ s^−1^ light and 70% relative humidity (RH). For gas-exchange experiments, 24-30-day-old plants were used.

Stomatal conductance of intact plants was measured using a rapid-response gas exchange measurement device consisting of eight through-flow whole-rosette cuvettes [39]. Prior to the experiment, plants were acclimated in the measurement cuvettes in ambient CO_2_ concentration (∼400 ppm), 100 µmol m^−2^ s^−1^ light (if not stated otherwise) and ambient humidity (RH 65-80%) for at least 1 h until stomatal conductance was stable. Thereafter, the following stimuli were applied: decrease/increase in CO_2_ concentration, darkness, reduced air humidity, and ozone. CO_2_ concentration was decreased to 100 ppm by filtering air through a column of granular potassium hydroxide. In CO_2_ enrichment experiments, CO_2_ was increased by adding it to the air inlet to achieve a concentration of 800 ppm. Darkness was applied by covering the measurement cuvettes. In blue-light experiments, dark-adapted plants were exposed to blue light (50 µmol m^−2^ s^−1^) from a LED light source (B42180, Seoul Semiconductor, Ansan, South Korea). The decreased/increased CO_2_ concentration, darkness and blue light were applied for 58 minutes. In the long-term elevated CO_2_ experiment (Fig 1D and S1E Fig), CO_2_ concentration was increased from 400 ppm to 800 ppm for 2.5 hours. To calculate stomatal half-response times, the whole 2.5-hour stomatal response to elevated CO_2_ was scaled to a range from 0 to 100% and the time when 50% of stomatal closure had occurred was calculated. Humidity was decreased by a thermostat system to 30-40% RH and stomatal conductance was monitored for another 56 min. In ozone experiments the plants were exposed to 350-450 ppb of ozone for 3 minutes and stomatal conductance was measured for 60 minutes after the start of the exposure.

In ABA-induced stomatal closure experiments, 5 µM ABA solution was applied by spraying as described in [40]. At time point 0, plants were removed from cuvettes and sprayed with either 5 µM ABA solution (5 μM ABA, 0.012% Silwet L-77 (PhytoTechnology Laboratories) and 0.05% ethanol) or control solution (0.012% Silwet L-77 and 0.05% ethanol). Thereafter plants were returned to the cuvettes and stomatal conductance was monitored for 56 minutes.

In ABA-induced inhibition of stomatal opening experiments, plants were acclimated in measurement cuvettes in darkness. At time point 0, plants were removed from cuvettes and sprayed with 2.5 µM ABA solution (2.5 μM ABA, 0.012% Silwet L-77 (PhytoTechnology Laboratories) and 0.05% ethanol) or control solution (0.012% Silwet L-77 and 0.05% ethanol). Thereafter plants were returned to the cuvettes, dark covers were removed and stomatal conductance was monitored in light for 56 minutes.

Prior to the measurement of diurnal pattern of stomatal conductance, plants were pre-incubated in the measurement cuvette at least 12 h in respective light and humidity conditions. Plants were measured in 16-minute intervals. Water Use Efficiency (WUE) was calculated based on the data of diurnal experiments as an average of day-time light period (from 9:00 to 17:00).

CO_2_-induced stomatal conductance in S2 Fig. Five-week old healthy plants, growing in a growth chamber with 70% humidity and 16 h light/8 h dark light condition, were used for stomatal conductance analyses at different CO_2_ concentrations by a LiCOR-6400XT as previously described (9). Relative stomatal conductance values were normalized relative to the average of ten data points preceding the [CO_2_] transitions (400 to 800 or 1000 ppm). The data presented are means of at least 4 leaves per genotype ± SEM.

### Stomatal aperture

The *MPK12* deletion mutant *mpk12-4* and wild type plants were grown in a growth chamber at 70% humidity, 75 μmol^m-2 s-1^ light intensity, 21°C, and 16 h light/8 h dark regime. Two-week old leaf epidermal layers of both genotypes were preincubated in opening buffer (10 mM MES, 10 mM KCl, and 50 mM CaCl_2_ at pH 6.15) for 2 h and individually stomata were imaged and tracked for measurement as before treatment. After that the leaf epidermal layers were incubated with buffers containing 10 μM ABA for 30 min and the individually tracked stomata were imaged. Stomatal apertures were measured by ImageJ software and genotype-blind analyses were used. The data presented are means and SEM n=3 experiments, 30 stomata per experiment and condition.

### Stomatal index and density

Plants at the age of 28-30 days were used for stomatal index and density measurements. Rosette leaves of equal size were excised and the abaxial side was covered with the dental resin (Xantopren M mucosa, Heraeus Kulzer, Germany). Transparent nail varnish was applied onto the dried impressions after the removal of the leaves. The hardened nail varnish imprints were attached onto a microscope glass slide with a transparent tape and imaged under Zeiss SteREO Discovery.V20 stereomicroscope. For quantification an image with the coverage of 0.12 mm^2^ was taken from the middle of the leaf, next to the middle vein. In total 81-84 plants per line from 2 independent biological repeats were analysed, one leaf from each plant, one image from each leaf. Stomatal index was calculated with the following formula: SI=Stomatal density/(Density of other epidermal cells + Stomatal density).

### Stomatal complex length

For the stomatal complex length measurements plants at the age of 28-35 days were used. Whole leaves were pre-incubated for 4 h abaxial side down in the buffer (10 mM MES, 5 mM KCl, 50 µM CaCl_2_, pH (with TRIS)) in the light. 4-6 plants per genotype and one leaf per plant were analyzed, altogether 84-126 stomatal complexes per genotype were measured.

### Y2H interaction tests

Interactions between MPK12 and selected protein kinases and phosphatases were tested in pairwise split-ubiquitin Y2H assays using the DUALhunter and DUALmembrane 3 kits (Dualsystems Biotech). For bait construction, the coding sequences of *MPK12* were PCR-amplified from total cDNAs from Col-0 and Cvi-0. Other *MPK12* variants with point mutations (K70R, Y122C and D196G+E200A) were created by two-step overlap PCR using the Col-0 *MPK12* as a template. *HT1* was also PCR-amplified from Col-0 cDNA. All *MPK12*s and *HT1* were digested with SfiI and cloned to the corresponding site in pDHB1, containing the Cub-LexA-VP16 fusion. For prey constructs, coding sequences of each selected gene was amplified from total Col-0 cDNAs, digested with SfiI, and cloned into either pPR3-N ( *HT1, OST1, BLUS1, IBR5, MKP2, MPK12, MPK12G53R, MPK11*) or pPR3-STE (*SnRK2.2, SnRK3.11, ABI1, ABI2, HAB1, HAB2*), containing a mutated NubG. All primers used are listed in Table S1. The pAI-Alg5 with a native NubI was used as a positive prey control, whereas the pDL2-Alg5 containing NubG served as a negative control.

For pairwise Y2H assays, the yeast strain NMY51 was co-transformed with bait and prey plasmids, and grown on SD-Leu-Trp plates to select for presence of both plasmids. At least 10 colonies from each transformation were pooled and resuspended in water to an OD600 of 0.5 from which 100, 1000, and 10000x serial dilutions were prepared, and spotted on SD-Leu-Trp and SD-Leu-Trp-His-Ade plates. SD-Leu-Trp plates were incubated at 30°C for 2 d, photographed and used for β-galatosidase overlay assays. SD-Leu-Trp-His-Ade plates were incubated for 2-4 d and photographed. The quantitative-β-galactosidase assay was performed with three pools of 10 independent colonies from each pairwise combination using the Yeast β-galactosidase assay kit (Thermo Scientific) by the non-stop quantitative method.

### BiFC interaction

Binary constructs containing split YFPs were designed and generated for cloning genes of interest by the ligation independent cloning (LIC) method. First, YFPn (amino acids 1-173 of eYFP), YFPc (amino acids 155-279 of eYFP) and the full-length YFP were amplified by multi-PCR steps to incorporate sequences for LIC method and the HA tag at 5’ and 3’ end, respectively. The PCR products were digested by EcoRI and cloned into the modified p35S/pCAMBIA1390 at the EcoRI/PmlI site to create 35S:YFPn, and 35S:YFPc in the pCAMBIA1390 vector, respectively.

For subsequent cloning, each gene of interest was amplified by two consecutive PCR reactions: first with gene-specific primers, and later with a pair of universal primers designed specifically for the LIC method. All primers used are listed in Table S1. To prepare vectors for LIC, plasmids of 35S:YFPn and 35S:YFPc were linearized by PmlI digestion, followed by T4 DNA polymerase treatment with dGTP to create 15-16 nucleotide 5’-overhangs. For insert preparation, the final PCR products of target genes were incubated with T4 DNA polymerase in the presence of dCTP to create the complementary overhangs with the vectors. Both vector and insert were mixed at room temperature, and proceeded with *E. coli* transformation after 5 min. The final constructs were sequence verified, and transformed to *Agrobacterium tumefaciens* GV1301 for agro-infiltration experiments.

For the BiFC assays, three different agrobacterial clones each harboring a YFPn fusion, a YFPc fusion, or the gene silencing suppressor P19 were co-infiltrated to the leaves of *Nicotiana benthamiana* at an OD600 of 0.2 for each clone in the infiltration buffer (10mM MES, 10mM MgCl_2_, 200μM acetosyringone). Images were acquired at 3 dpi with a Zeiss LSM710 confocal microscope. The YFP signals were excited by a 514nm laser, and emission between 518-564 nm was collected. All of the acquisition parameters were kept identical within each experiment, and only images acquired from the same leaves with multiple infiltration spots were compared.

### Split luciferase complementation assay

The *MPK12* cDNA was cloned into a vector containing the N-439 terminal half of luciferase (nLUC) and *HT1* was cloned into the cLUC. The constructs in the Agrobacterium strain GV3101 were co-infiltrated into *N. benthamiana* leaves with P19 at an OD600 of 0.8. The infiltrated leaves after three days of infiltration were harvested for bioluminescence detection. Images were captured with a CCD camera.

### Measurement of S-type anion currents

*Arabidopsis* guard cell protoplasts were isolated as described previously [41]. Guard cell protoplasts were washed twice with washing solution containing 1 mM MgCl_2_, 1 mM CaCl_2_, 5 mM MES and 500 mM D-sorbitol (pH 5.6 with Tris). During patch clamp recordings of S-type anion currents, the membrane voltage started at +35 to −145 mV for 7 s with −30 mV decrements and the holding potential was +30 mV. The bath solutions contained 30 mM CsCl, 2 mM MgCl_2_, 10 mM MES (Tris, pH 5.6), and 1 mM CaCl_2_, with an osmolality of 485 mmol/kg. The pipette solutions contained 5.86 mM CaCl_2_, 6.7 mM EGTA, 2 mM MgCl_2_, 10 mM Hepes-Tris (pH 7.1), and 150 mM CsCl, with an osmolality of 500 mmol/kg. The free calcium concentration was 2 µM. The final osmolalities in both bath and pipette solutions were adjusted with D-sorbitol. Mg-ATP (5 mM) was added to the pipette solution before use. 13.5 mM CsHCO_3_ (11.5 mM free [HCO_3_^-^] and 2 mM free [CO_2_]) was freshly dissolved in the pipette solution before patch clamp experiments. The concentrations of free bicarbonate and free CO_2_ were calculated using the Henderson–Hasselbalch equation (pH = pK1+log [HCO_3_^-^]/[CO_2_]). pK1 = 6.352 was used for the calculation. [HCO_3_^-^] represents the free bicarbonate concentration and [CO_2_] represents the free CO_2_ concentration.

### Protein expression and purification

For *in vitro* kinase assays, HT1, HT1 K113M, MPK11, MPK12, MPK12 G53R, MPK12 K70R and MPK12 Y122C were cloned into pET28a vector (Novagen, Merck Millipore) using primers listed in S1 Table. Point mutations corresponding to K113M in HT1, K70R in MPK12 and Y122C in MPK12 were created with two-step PCR using primers listed in S1 Table.

6xHis-HT1WT, 6xHis-HT1 K113M, 6xHis-MPK12 WT, 6xHis-MPK12 G53R, 6xHis-MPK12 K70R, 6xHis-MPK12 Y122C and 6xHis-MPK11 WT were expressed in *E. coli* BL21(DE3) cells. A 2 mL aliquot of an overnight culture was transferred to a fresh 1 L 2xYT medium and grown at 37 °C to an absorbance of ∼0,6 at OD600. The cultures were chilled to 16 °C and recombinant protein expression was induced by 0.3 mM isopropyl b-D-thiogalactopyranoside for 16 h. The cells were harvested by centrifugation (5000 rpm, 10 min, 4 °C) and stored at −80 °C until use.

All purification procedures were carried out at 4 °C. The cells were resuspended in 30 mL of lysis buffer (50 mM Tris-HCl (pH 7.4), 300 mM NaCl, 5% (v/v) glycerol, 1% (v/v) Triton X-100, 1 mM PMSF, 1 μg/ml aprotinin, 1 μg/ml pepstatin A, 1 μg/ml leupeptin) and lysed using an Emulsiflex C3 Homogenizer. Cell debris was removed by centrifugation at 20 000 rpm for 30 min. The protein-containing supernatant was mixed for 1 h at 4 °C with 0.20 mL of Chelating Sepharose Fast Flow resin (GE Healthcare), charged with 200 mM NiSO_4_ and pre-equilibrated in the lysis buffer. The protein-resin complex was packed into a column and the beads were washed with 5x10 column volumes (CV) of a wash buffer I (50 mM Tris-HCl (pH 7.4), 600 mM NaCl, 5% (v/v) glycerol, 1% (v/v) Triton X-100), 5x10 CV of a wash buffer II (50 mM Tris-HCl (pH 7.4), 300 mM NaCl, 5% (v/v) glycerol, 0.1% (v/v) NP-40) and 2x10 CV of a wash buffer III (50 mM Tris-HCl (pH 7.4), 150 mM NaCl, 5% (v/v) glycerol, 0.1% (v/v) NP-40). The protein was eluted by incubating the beads for 5 min at room temperature with an imidazole containing elution buffer (50 mM Tris-HCl, 150 mM NaCl, 5% (v/v) glycerol, 0.1% (v/v) NP-40, 300 mM imidazole). MPK12 proteins were concentrated and imidazole was removed by Millipore Amicon Ultra-0.5 Centrifugal Filter Concentrators (NMWL 3000). Glycerol was added to a final concentration of 20% (v/v) and 20 μL aliquots of the eluted protein were snap frozen in liquid nitrogen and stored at −80 °C.

### *In vitro* kinase assays

Protein concentrations were estimated on 10% SDS-polyacrylamide gel using BSA as a standard. HT1 kinase activity assay was performed by incubating constant amount of purified recombinant HT1 and 0-30 μM MPK12 or 0-10 μM MPK11 in a reaction buffer (50 mM Tris-HCl (pH 7.4), 150 mM NaCl, 20 mM MgCl2, 60 mM imidazole, 1 mM DTT, 0.2 mg/ml insulin) at room temperature for 10 min. Then casein (1 mg/ml), 500 μM ATP and 100 μCi/ml 32P-γ-ATP were added and reaction aliquots were taken at 30 min time point. Reactions were stopped by the addition of SDS loading buffer. Proteins were separated on a 10% SDS-polyacrylamide gel and visualized by Coomassie brilliant blue R-250 (Sigma) staining. HT1 activity was determined by autoradiography and quantified by ImageQuant TL Software.

### Model of MPK12

Protein structure homology modelling of MPK12 was done with SWISS-MODEL [42] using human MPK7 as a template and visualized with Jmol (http://jmol.sourceforge.net/).

### Phylogeny analysis

Profile hidden Markov model (HMM) of MPKs was built with hmmbuild of HMMR3.1b1 using all 20 Arabidopsis thaliana MPKs' full protein sequences aligned with ClustalW. The protein sequences from other analyzed species was downloaded from Phytozome v10 (proteins, primary transcript only) and searched for MPK HMM with hmmsearch of HMMR3.1b. The full-length protein sequences from all species matching MPK HMM were aligned with Muscle and phylogeny tree was generated using PhyML with following settings: Model=LG, Tree searching operations=best of NNI & SPR, Starting tree=BioNJ, Number of bootstrapped data sets = 1000. Three Cyclin-dependent kinases were included in the analysis as an outgroup. Another phylogram was drawn from the same protein sequences using PASTA with similar results (not shown). The protein sequences from other species were named according to the closest relative found in Arabidopsis thaliana using TAIR BLAST 2.2.8. A FASTA file will all protein sequences is provided as S2 Table.

### Statistical analysis

Statistical analyses were performed with Statistica, version 7.1 (StatSoft Inc., Tulsa, OK, USA). All effects were considered significant at p < 0.05.

## Acknowledgments

Tuomas Puukko provided excellent technical assistance. We thank Aleksia Vaattovaara for assistance with the phylogeny analysis.

